# Targeted Deletion of *Fgf9* in Tendon Disrupts Mineralization of the Developing Enthesis

**DOI:** 10.1101/2022.08.25.505295

**Authors:** Elahe Ganji, Connor Leek, William Duncan, Debabrata Patra, David M. Ornitz, Megan L. Killian

## Abstract

The enthesis is a transitional tissue between tendon and bone that matures postnatally. The development and maturation of the enthesis involve cellular processes likened to an arrested growth plate. In this study, we explored the role of fibroblast growth factor 9 (*Fgf9*), a known regulator of chondrogenesis and vascularization during bone development, on the structure and function of the postnatal enthesis. First, we confirmed spatial expression of *Fgf9* in wildtype tendon and enthesis using *in situ* hybridization. We then used Cre recombinase driven by the scleraxis promoter (ScxCre) to conditionally inactivate *Fgf9* in mouse tendon and enthesis. Characterization of enthesis morphology and mechanical properties in *Fgf9*^*ScxCre*^ and wildtype (WT) entheses showed a smaller calcaneal and humeral apophyses, thinner cortical bone at the attachment, increased cellularity, and reduced failure load in mature entheses in *Fgf9*^*ScxCre*^ compared to WT littermates. During postnatal development, we found reduced chondrocyte hypertrophy and disrupted type X collagen (Col X) in *Fgf9*^*ScxCre*^ entheses. These findings support a model in which tendon-derived *Fgf9* regulates the functional development of the enthesis, including its postnatal mineralization.

## Introduction

The tendon-bone attachment (enthesis) is critical for the transmission of muscle-generated loads to the vertebrate skeleton. The enthesis forms embryonically as a compliant anchorage between tendon and bone.^1,2^ The fibrocartilage enthesis matures postnatally into a graded transitional tissue with increasing mineral and proteoglycan content that reinforces the fibrous tendon into mineralized bone,^3^ and its morphology mimics that of an arrested growth plate.^4–6^ Growth plates of long bones and fibrocartilage entheses form primarily via endochondral ossification. Additionally, the cellular patterns of the developing enthesis form from a pool of progenitor cells that express *Sox9*, which is also a major regulator of the growth plate in long bones. However, the development of these two structures is not identical, as the growth plate fuses with age (∼2-8 months of age in mice, depending on anatomical site), yet the enthesis remains fibrocartilaginous throughout the lifespan. In addition, the *Sox9*+ progenitor cells that establish the enthesis also co-express *Scleraxis*, which is not expressed in the growth plate.^7,8^ Therefore, a gap in knowledge exists in identifying the similar and divergent patterns of the developing enthesis and growth plate during postnatal growth.

Recent studies by our laboratory and others have identified the potential role of fibroblast growth factors (*FGFs)* in the formation and adaptation of the entheses.^9,10^ Several FGF ligands and their binding receptors (FGFR) are critical for growth plate development.^11–18^ Specifically, FGF18 regulates chondrocyte proliferation and differentiation of the growth plate during bone development.^15^ FGF9, together with FGF2 and 18, can compensate for each other during bone growth.^12,19–21^ Spatially, FGF9 is most prevalent in the perichondrium and periosteum^22^ and *Fgf9* expression regulates chondrocyte proliferation and hypertrophy through its affinity to FGFR3.^22^ We and others have shown that global deletion of *Fgf9* during embryonic bone development results in reduced chondrocyte proliferation, delayed hypertrophy, and limb shortening in mouse embryos.^9^ Global deletion of *Fgf9* in mouse embryos also leads to enlarged tuberosities, which are sites of tendon entheses.^9,12,21^ Despite its prominent role in bone growth, the role of *Fgf9* during enthesis development remains unknown.

In this study, we aimed to identify the role of *Fgf9* in *Scx*-lineage cells, as Scx is an early marker of the tendon/ligament progenitors, and its expression is essential for the formation and postnatal growth of the enthesis.^23–27^ *Scx*-positive chondroprogenitors also contribute to chondrocyte differentiation at the bony eminence of the enthesis.^28,29^ We generated mice to conditionally inactivate *Fgf9* in Scx-lineage cells (using ScxCre) to study the structural and functional role of FGF9 on the postnatal development of the fibrocartilage enthesis. We compared the mineral and cellular morphology as well as functional (mechanical) properties of the mature fibrocartilage entheses for both Achilles and supraspinatus attachments between normally developing (wildtype) and *Fgf9*^*ScxCre*^ mice. We hypothesized that *Fgf9*^*ScxCre*^ mice would develop disruptions in postnatal mineralization and organization of the mature fibrocartilaginous enthesis with impaired mechanical properties and reduced mineral content in the apophysis compared to normally developing littermates.

## RESULTS

### *Fgf9* is expressed in the Achilles tendon and attachment and loss of *Fgf9* in tendon progenitors leads to impaired apophyseal and entheseal growth

*Fgf9* is robustly expressed postnatally (postnatal day 0) in the neonatal enthesis and tendon (Figure 1). FGF9 deletion in the tendon progenitors led to reduced trabecular bone volume and trabecular number at the humeral epiphysis compared to WT controls, as measured using microCT (Table 1). The calcaneus of *Fgf9*^*ScxCre*^ mice was significantly shorter and had reduced bone and tissue volume compared to WT (Table 1). *Fgf9*^*ScxCre*^ mice had lower tissue volume in the adult humeral and calcaneal epiphyses (Figure 2) and thinner subchondral bone thickness at supraspinatus attachment at the humeral head and Achilles tendon attachment on the calcaneus (Figure 2).

**Table 1:** Bone morphometric measurements at the site of Achilles and supraspinatus attachments at 8wk of age. (Mean ± standard deviation; p<0.05)

| Anatomical Site | WT | *Fgf9*<sup>ScxCre</sup> | *p*-value |
| --- | --- | --- | --- |
| **Humerus** |  |  |  |
| Length (mm) | 11.76 ± 0.55 | 11.31 ± 0.56 | 0.0919 |
| Cortical thickness at mid-diaphysis (mm) | 0.17 ± 0.02 | 0.15 ± 0.02 | 0.2870 |
| Cortical BMD at mid-diaphysis (g/cm<sup>3</sup>) | 0.73 ± 0.03 | 0.757 ± 0.04 | 0.1295 |
| **Humeral epiphysis** |  |  |  |
| TV (mm<sup>3</sup>) | 2.42 ± 0.17 | 2.14 ± 0.13 | 0.0045* |
| BV (mm<sup>3</sup>) | 0.95 ± 0.17 | 0.87 ± 0.16 | 0.2135 |
| BV/TV (%) | 39.34 ± 6.23 | 40.78 ± 7.14 | 0.3592 |
| Trabecular TV (mm<sup>3</sup>) | 1.21 ± 0.15 | 1.13 ± 0.09 | 0.1609 |
| Trabecular BV (mm<sup>3</sup>) | 0.42 ± 0.08 | 0.35 ± 0.06 | 0.0431* |
| Trabecular BV/TV (%) | 35.22 ± 5.34 | 30.40 ± 4.23 | 0.0572 |
| Trabecular number (1/mm) | 5.19 ± 0.25 | 4.83 ± 0.30 | 0.0254* |
| Trabecular thickness (μm) | 0.07 ± 0.01 | 0.06 ± 0.01 | 0.1367 |
| Trabecular separation (μm) | 0.17 ± 0.02 | 0.17 ± 0.020 | 0.4046 |
| **Calcaneus** |  |  |  |
| Length (mm) | 3.75 ± 0.13 | 3.62 ± 0.12 | 0.0431* |
| TV (mm<sup>3</sup>) | 2.80 ± 0.39 | 2.36 ± 0.31 | 0.0175* |
| BV (mm<sup>3</sup>) | 2.08 ± 0.45 | 1.59 ± 0.37 | 0.0239* |
| BV/TV (%) | 73.55 ± 7.07 | 66.97 ± 6.69 | 0.0495* |
| Cortical thickness (mm) | 0.20 ± 0.04 | 0.17 ± 0.04 | 0.0942 |
| Cortical BMD (g/cm<sup>3</sup>) | 0.76 ± 0.15 | 0.73 ± 0.16 | 0.6599 |
| **Calcaneal Epiphysis** |  |  |  |
| TV (mm<sup>3</sup>) | 0.14 ± 0.04 | 0.11 ± 0.03 | 0.0415* |
| BV (mm<sup>3</sup>) | 0.13 ± 0.04 | 0.09 ± 0.03 | 0.0644 |
| BV/TV (%) | 84.48 ± 6.02 | 82.75 ± 6.82 | 0.6241 |
| TMD (g/cm<sup>3</sup>) | 0.63 ± 0.10 | 0.54 ± 0.12 | 0.1825 |

**Figure 1:**
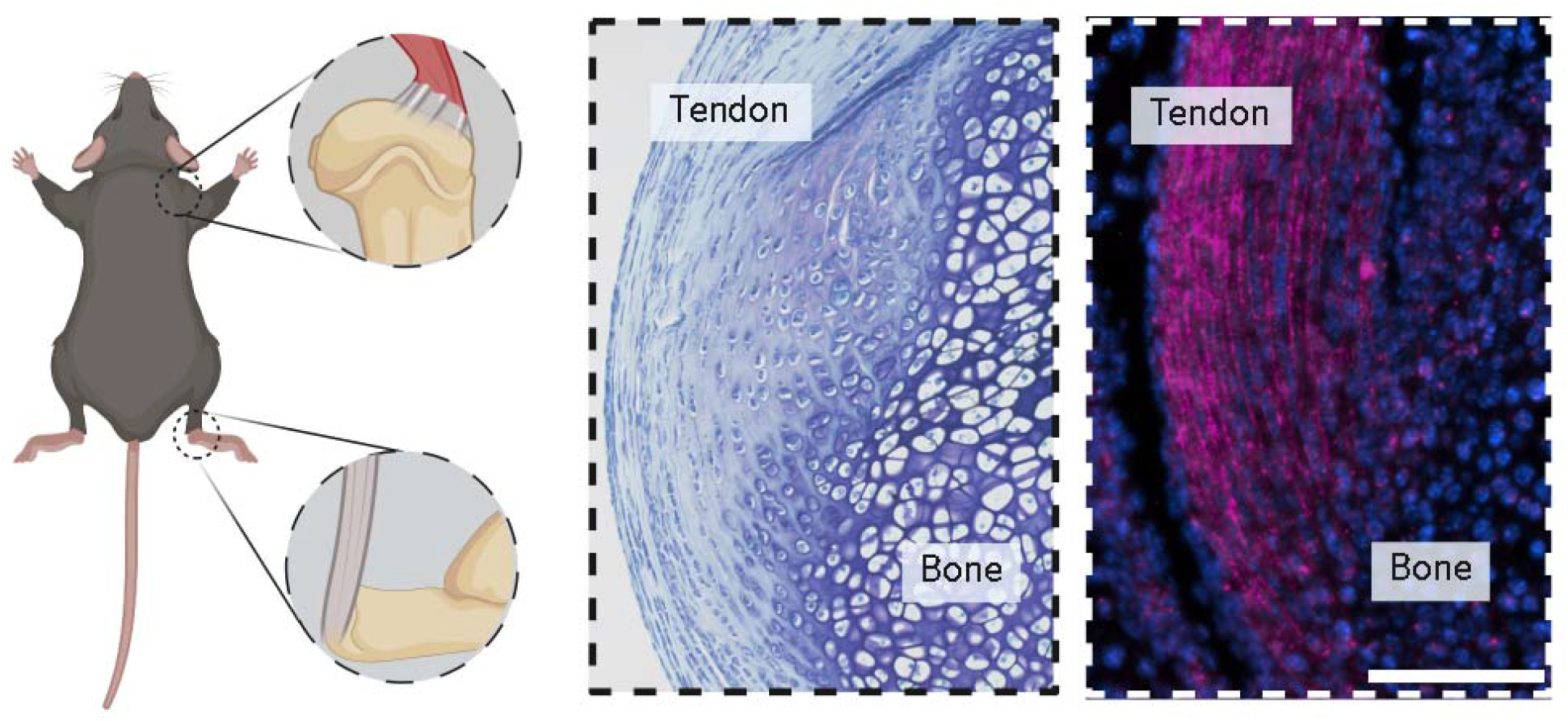
Entheses were analyzed from the supraspinatus tendon of the rotator cuff (shoulder) and the Achilles tendon (ankle) of mice. Resident cells of the wildtype Achilles tendon and enthesis, shown histologically with Toluidine Blue staining (middle panel), expressed elevated levels of *Fgf9* (magenta, right panel) compared to bone. Fluorescent *in situ* hybridization for Mm-*Fgf9*. Scale bar = 100 µm.

**Figure 2.**
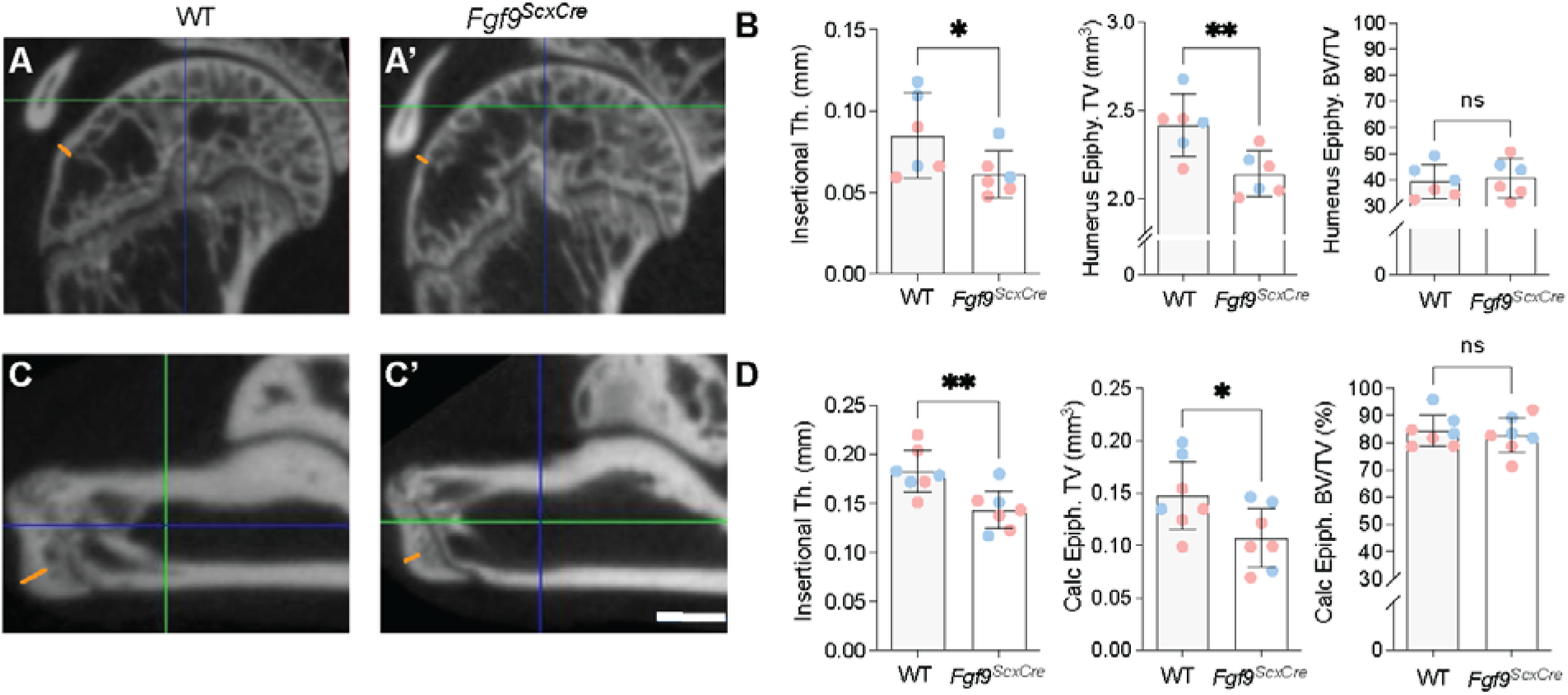
At 8 weeks of age, *Fgf9*^ScxCre^ mice had reduced insertional thickness (Th.) and tissue volume (TV) at the (B) humeral epiphyses and (D) calcaneal apophyses, but no difference in epi/apophyseal bone volume ratio (BV/TV), compared to age-matched WT littermates. Orange lines in panels A, A’, C, and C’ indicate entheses regions for insertional thickness measurements, with blue and green lines representing frontal/sagittal and transverse planes, respectively. Blue and pink dots in panels B and D denote male and female mice, respectively. Data presented as mean ± 95% CI; * = p<0.05; ** = p<0.01. Scale bar = 600 µm.

Cortical thinning of both the supraspinatus and Achilles entheses in *Fgf9*^*ScxCre*^ attachments was confirmed using histology (Figure 3). Additionally, *Fgf9*^*ScxCre*^ mice developed an acellular metachromatic region at the superior Achilles enthesis as indicated by lighter, pink staining at the insertion (Figure 3). In contrast, acellular regions of fibrocartilage were not evident in attachments of WT mice (Figure 3).

**Figure 3.**
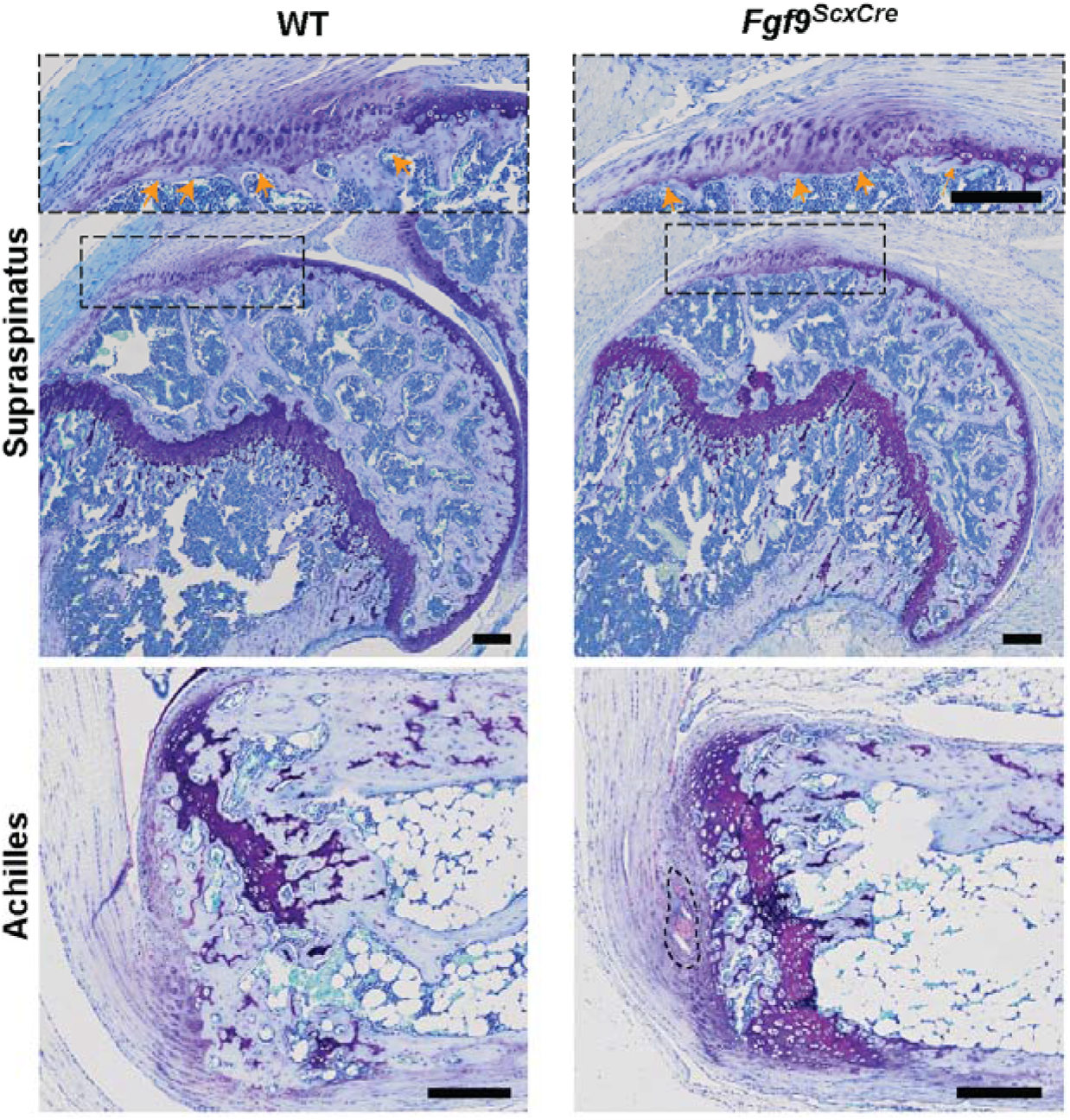
The supraspinatus enthesis of 8wk old *Fgf9*^*ScxCre*^ mice had thinner cortical bone compared to age-matched WT mice (orange arrowheads). *Fgf9*^*ScxCre*^ mice also had acellular metachromatic regions at the superior Achilles enthesis, shown in Toluidine blue stained sections of Achilles (outlined with dashed black line), as well as delayed calcaneal growth plate fusion compared to WT mice; scale bar = 100 µm scale.

### *Fgf9* deletion in the tendon progenitors resulted in increased cellularity and delayed maturation of the enthesis

During normal postnatal growth, the area of the Achilles enthesis (outlined in white dashed regions, Figure 4) increased from developing (0.03±0.003mm^2^) to young-adult samples (0.06± 0.014 mm^2^). Cellular density of the enthesis also decreased during postnatal development (Developing vs. Young-adult: p<0.0001; Developing vs. Adult: p<0.0001; Young-adult vs. Adult: p=0.0007) (Figure 4A’). In *Fgf9*^*ScxCre*^ mice, cellular density of adult entheses was higher at the supraspinatus (p < 0.01) and Achilles (p=0.0563) tendon attachment sites compared to WT mice (Figure 5).

**Figure 4.**
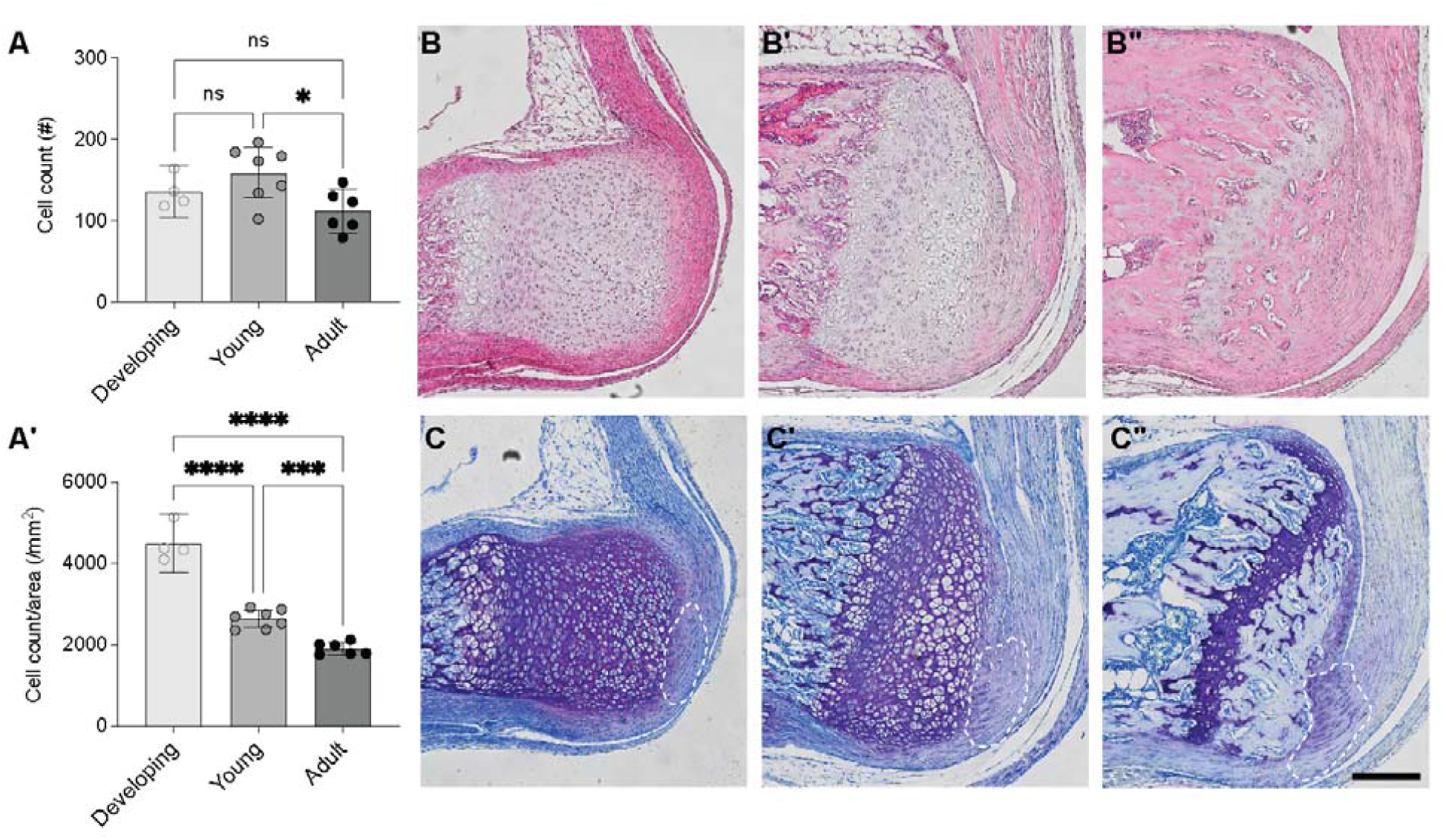
During postnatal development, the cellularity of the enthesis decreased. (A) Total cell count and (A’) cell count per area (cellularity) were measured using ImageJ. Histological representation of the Achilles enthesis stained using (B) H&E and (C) Toluidine Blue during developing stages, as well as for the (B’, C’) young-adult enthesis and (B”, C”) adult enthesis. Enthesis area outlined in white dashed regions. High magnification insets show the tendon attachment sites (arrow) that we quantified in A, A’. Scale bar = 200 µm. Data presented as mean ± 95% CI; * = and * = p<0.05; *** = p<0.0007; **** = p<0.0001.

**Figure 5.**
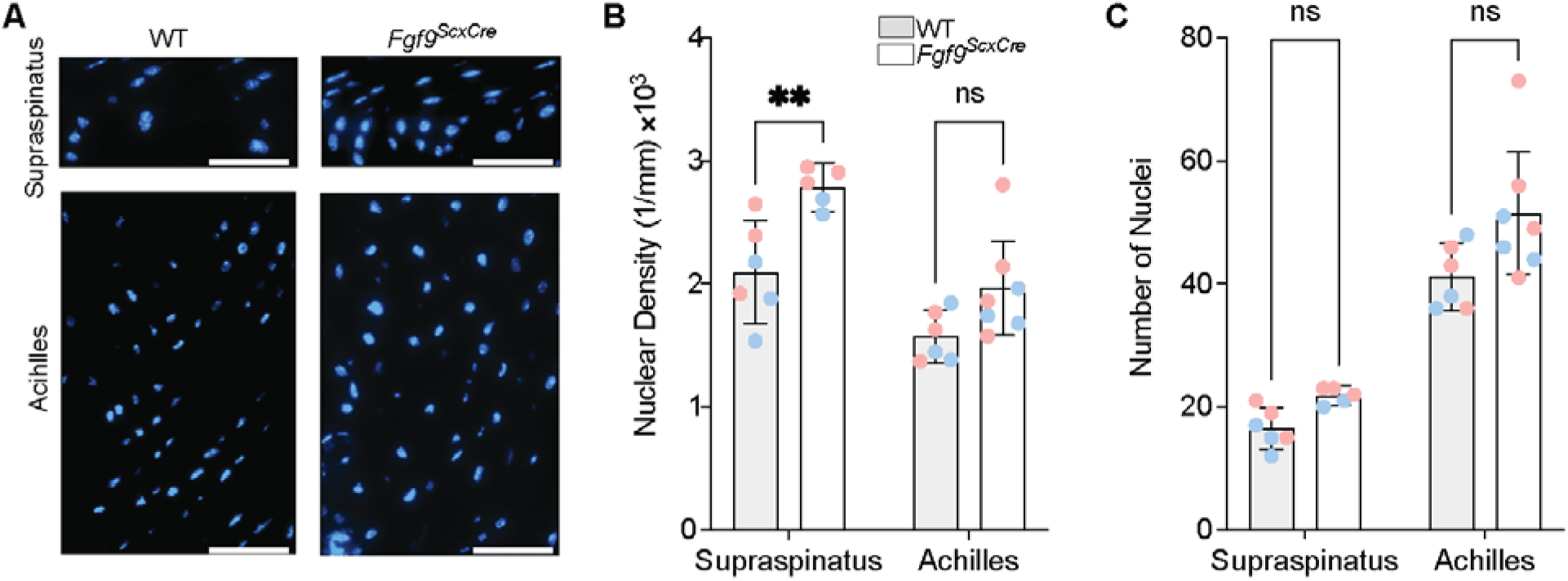
Cellular density was higher in supraspinatus entheses of *Fgf9*^*ScxCre*^ mice compared to WT. (A) Representative fluorescent images of supraspinatus and Achilles’ regions of interest (scale bar 20 µm). (B) Nuclear density and (C) number of nuclei presented for both *Fgf9*^*ScxCre*^ and WT mice at 8wk of age; Pink dots/lines = female mice; Blue dots/lines = male mice. Scale bar = 50 µm. Data presented as mean ± 95% CI and ** = p < 0.01.

### Tendon/enthesis-specific deletion of *Fgf9* led to impaired enthesis mechanical properties

To measure the mechanical properties of the enthesis in the presence or absence of FGF9, we uniaxially loaded the dissected tendon/enthesis complex in a heated water bath under tensile tension. Achilles tendon entheses of *Fgf9*^*ScxCre*^ mice had reduced ultimate load at 8wk of age (p= 0.041, Figure 6A,C) with no change in cross-sectional area (CSA; Figure 6B, p=0.77), stiffness (p=0.14, Table 2), or work to max load (p=0.11, Table 2). Tensile mechanical properties did not change between *Fgf9*^*ScxCre*^ and WT tendon entheses (elastic modulus: p= 0.64, Table 2; maximum stress: p=0.17, Table 2; strain at max stress: p= 0.32, Table 2; or toughness: p=0.12, Table 2).

**Table 2:** Biomechanical properties of the Achilles attachments at 8wk of age (mean ± standard deviation; unpaired t-tests).

|  | WT | *Fgf9<sup>ScxCre</sup>* | *p*-value |
| --- | --- | --- | --- |
| Young's modulus (MPa) | 138.4 ± 37.7 | 124.2 ± 57.0 | 0.645 |
| Stiffness (N/mm) | 11.23 ± 2.11 | 9.09 ± 2.26 | 0.142 |
| Work to max. load (mJ) | 4.56 ± 0.57 | 3.26 ± 1.55 | 0.112 |
| Ultimate stress (N/mm<sup>2</sup>) | 25.36 ± 5.67 | 19.31 ± 7.42 | 0.170 |
| Ultimate strain (mm/mm) | 298.5 ± 27.5 | 264.7 ± 66.5 | 0.318 |
| Toughness (KJ/mm<sup>2</sup>) | 3,372 ± 924 | 2,387 ± 985 | 0.124 |

**Figure 6.**
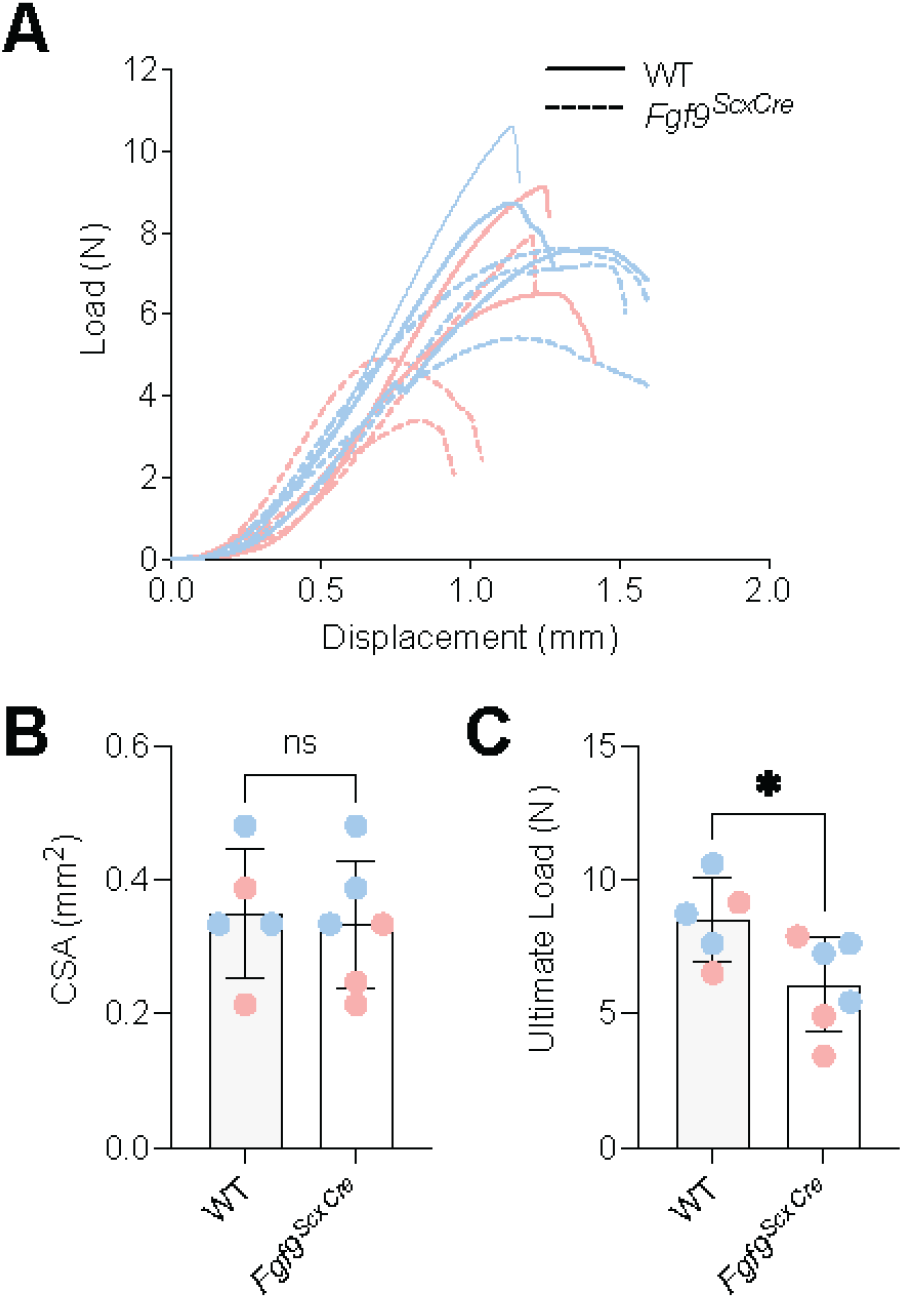
Achilles tendon entheses of 8wk old *Fgf9*^*ScxCre*^ mice had reduced ultimate load compared to age-matched WT mice. (A) Overlayed load-displacement curves, (B) cross-sectional area (CSA), and, (C) maximum load for all samples tested. Pink dots/lines = female mice; Blue dots/lines = male mice. Error bars denote mean ± 95% CI, * = p<0.05; ns= not significantly different.

### Loss of *Fgf9* in enthesis progenitors resulted in delayed mineralization

To investigate the underlying changes associated with structural adaptation in adult enthesis, the mineralization of the secondary ossification center (SOC) was characterized using Masson’s Trichrome staining and immunohistochemistry (IHC) for type X collagen (Col X). Young *Fgf9*^*ScxCre*^ calcanei had fewer hypertrophic chondrocytes at the SOC compared to WT (Figure 7 A, B). Interestingly, while the size of the SOC did not significantly differ between groups (p=0.20, Figure 7C), the Col X+ area within the SOC was smaller in *Fgf9*^*ScxCre*^ mice compared to WT mice (p= 0.035; Figure 7 D-F).

**Figure 7.**
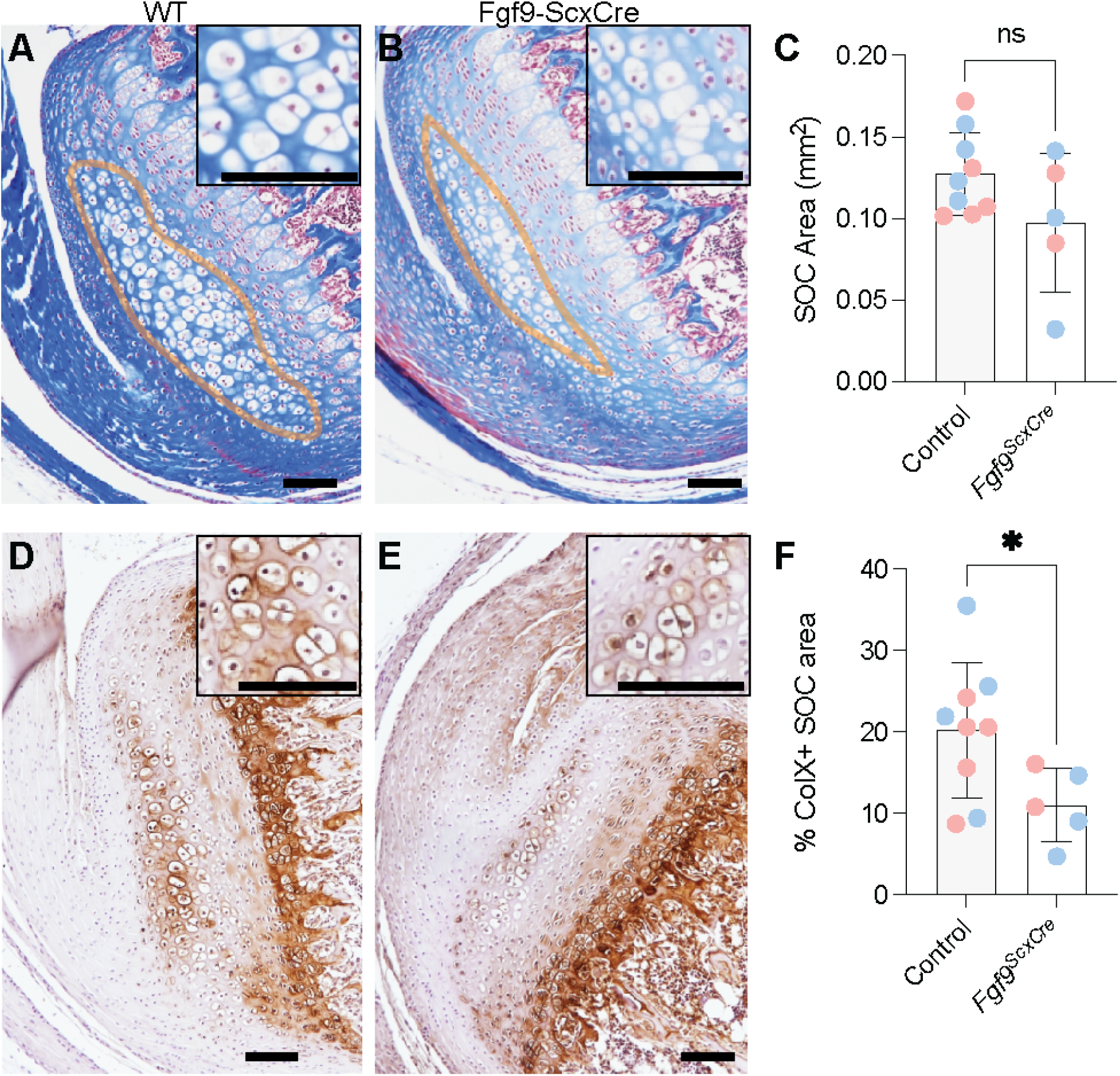
(A, B) 3-week-old *Fgf9*^*ScxCre*^ mice had smaller SOC areas (A, B; outlined in orange; quantified in C) and smaller ColX+ areas (D-F) compared to age-matched WT littermates. * = p < 0.05; ns = not significantly different (p > 0.05). Scale bar = 100µm. Blue dots in C and F indiciate male mice and pink indicates female mice.

## DISCUSSION

In this study, we investigated the role of tendon/enthesis-derived *Fgf9* on the maturation of two functionally graded, fibrocartilaginous entheses (i.e., Achilles and supraspinatus). We showed that, unlike the growth plate^30^, *Fgf9* is expressed postnatally in tendon and enthesis. We then showed that constitutive deletion of *Fgf9* using ScxCre led to smaller but more cellular tendon interfaces with fewer hypertrophic chondrocytes and less Col X+ matrix in the maturing apophysis. This suggests a potent disruption and delay in endochondral-like bone formation driven by loss of *Fgf9* in the enthesis and SOC, similar to findings from the growth plates of global *Fgf9*^*-/-*^ mutants.^21^ These mineralization delays may have contributed to the lower biomechanical properties (e.g., ultimate load) of the tendon enthesis.

Others have also shown that disruption of the mineral matrix can significantly reduce the tensile strength of the enthesis.^31,32^ Surprisingly, despite the thinner mineralized bone at the adult supraspinatus and Achilles entheses, *Fgf9*^*ScxCre*^ mice had increased cellular density at the attachment site. Previously, increased mesenchymal cellularity has been linked to the delayed palatal growth in embryos with global deletion of *Fgf9*.^33^ Thus, the increased cellularity we observed is likely due to delayed maturation in the *Fgf9*^ScxCre^ enthesis. Additionally, the acellular metachromatic defect at the Achilles enthesis of *Fgf9*^ScxCre^ mice may indicate an altered extracellular matrix at the tendon-bone attachment, caused by changes in compressive loading of the tendon that results in increased proteoglycan formation.^34–36^ In models of enthesis unloading, such as tenectomy or hindlimb unloading, markers of chondrogenic differentiation (e.g., PTHrP) are reduced at the tendon-bone interface.^37^ It is possible that cell differentiation and matrix production are mediated by interactions between FGF signaling and other major signaling pathways, such as Hedgehog signaling. Lastly, a potential explanation for the reduced bone morphometric properties may be linked to decreased effective load transfer through the supraspinatus enthesis that manifests from decreased remodeling and mineral deposition near the enthesis.^38,39^

FGF9 plays a major role in vascularization of many tissues, including bone during development^21^ and in muscle with reperfusion following ischemia.^40^ The fibrocartilage entheses investigated in the present study are not considered well-vascularized tissues, yet the potential delays in maturation we observed may still be caused by delays in vascularization especially to the SOC.^41^ Additionally, others have reported that loss of FGF9 during growth leads to delayed vascular invasion and, consequently, delayed initiation of chondrocyte hypertrophy,^21^ which we also observed in the SOC of *Fgf9*^*ScxCre*^ mice. Future studies should explore the role of FGF9 and vascularization during development of the fibrocartilaginous enthesis.^42^

This study is not without limitations. While we focused primarily on the role of FGF9 in structure and function of fibrocartilage entheses, we did not investigate migratory or fibrous entheses, like the deltoid tuberosity or the medial collateral ligament, respectively. The resident progenitor cells in these other types of entheses have more dynamic turnover during growth^43^ and may offer a different perspective on FGF signaling during enthesis growth. Additionally, use of inducible Cre drivers, such as Gli1-CreERT2^44^ or Scx-CreERT2^45^ would reduce the potential off-target effects of FGF9 knockout in cartilage and perichondrium, as the constitutive ScxCre also targets chondrogenic progenitors and other cell types.

## MATERIALS AND METHODS

### Animal models

This study was approved by the Institutional Animal Care and Use Committees (IACUC) at the University of Delaware and the University of Michigan (n=52 mice total). Mice were housed in 12 hours on/off light cycle housing and placed in same-sex cages with littermates after weaning. Food and water were provided for ad libitum. To generate conditional knockout (*Fgf9*^*ScxCre*^) mice, we crossed *Fgf9*^*flx/+*^; ScxCre females with *Fgf9*^*flx/flx*^ males. Offspring were genotyped using PCR (Transnetyx, Cordova, TN). Both male and female ScxCre; *Fgf9*^*flx/flx*^ (*Fgf9*^*ScxCre*^) and wildtype (WT; *Fgf9*^*flx/flx*^) littermates were euthanized at 3 weeks (Young; n=7/genotype) and 8 weeks (Adult; n=8 *Fgf9*^*ScxCre*^ and n=10 WT) of age using carbon dioxide asphyxiation and thoracotomy. Both male and female mice were used, and a minimum of n = 3 per sex were collected for each time point and assay. Normal development of the fibrocartilage enthesis (i.e., Achilles’ attachment) was assessed using an additional group of male WT mice at the following time points: Developing (postnatal day, P7-10, n = 4), Young adult (P15-28, n = 7), and Adult (P45-129, n = 6). A third cohort of WT mice (n = 3) were used for *in situ* hybridization at P0. Both hindlimbs and forelimbs were dissected at the time of euthanasia for imaging and contralateral hindlimbs were kept intact and stored at 4°C for uniaxial tensile testing.

### In situ *hybridization*

Spatial expression of *Fgf9* was visualized in the neonatal tendon enthesis using *in situ* hybridization (RNAscope Multiplex Fluorescent Reagent Kit v2, Advanced Cell Diagnostics, Hayward, CA, USA). P0 hindlimbs were decalcified in 14% EDTA for 2 weeks, paraffin-embedded, and sectioned at 7µm thickness. Sections were labeled with mm*Fgf9* probe and both positive controls (probes targeting housekeeping genes *Polr2A, Ppib*, and *Ubc*) and negative controls (*dapB*) were used. Nuclei were counterstained with DAPI and sections were mounted with Citifluor Antifade mounting medium (Electron Microscopy Science, Hatfield, PA, USA). Slides were imaged at 40X magnification using a fluorescent microscope (Lionheart FX, BioTek, Winooski, Vermont, USA).

### Microcomputed tomography

Limbs were dissected and skin removed at time of euthanasia and immediately fixed in 4% paraformaldehyde for 24-48hr. To assess morphological bone differences between *Fgf9*^*ScxCre*^ and WT mice, fixed limbs were wrapped in 70% ethanol-soaked gauze and scanned using micro-computed tomography (microCT; Skyscan 1276, Bruker, Belgium) with acquisition settings optimized for mouse limb imaging (0.5mm Aluminum filter, 10.6 μm voxel size, 50 kV voltage, 200 mA current, and 950 ms exposure time, 0.3° rotation step, and 360° scan). Humeral epiphyses and calcaneal apophyses were segmented based on the growth plate morphology (humeral epiphysis: superior to the growth plate; calcaneal apophysis: posterior to the growth plate) using CTAN software (Bruker, Belgium). Tissue volume (TV), bone volume (BV), and bone volume ratio (BV/TV, %) in the calcaneus as well as the humeral and calcaneal epiphyses were measured, and for both the humerus and the calcaneus, bone length, bone mineral density (BMD), and diaphyseal cortical thickness were measured following mineral density calibration. Mineralized insertional thickness (mm) was measured manually in the mid-sagittal plane for the humeral head and calcaneus at the supraspinatus and Achilles tendon entheses, respectively, using CTAn Software (Bruker, Belgium). Thickness measurements were repeated three times along the anatomical site of the attachment for both Achilles and supraspinatus entheses. Percent variation was calculated between at least 3 in-plane slices to test for repeatability of measurement and values were averaged for quantitative analysis (standard error of mean <3%).

### Histology

Fixed tissues were decalcified (Formical□2000, StatLab, McKinney, TX), paraffin embedded, and sectioned at 6 μm thickness. For staining, Toluidine Blue and Hematoxylin & Eosin (H&E) stains were used for qualitative assessment of proteoglycan and overall enthesis morphology at the Achilles and supraspinatus entheses, respectively. Additionally, sections were stained with Masson’s Trichrome to assess formation of the fibrocartilaginous enthesis. Stained slides were imaged using brightfield microscopy (Imager A2 microscope, Carl Zeiss, Germany).

Enthesis cellularity was measured from Toluidine Blue-stained slides of normally developing neonatal, young-adult, and adult samples. Entheses were manually segmented based on the cellular morphology and the change in GAG distribution, using particle analysis in ImageJ software.^46^ In the adult WT and *Fgf9*^ScxCre^ attachment sites, cell nuclei were stained with DAPI. Fluorescent images were taken using the DAPI channel (20x objective, Axio Observer.Z1 microscope, Carl Zeiss, Germany). The total number of nuclei in a field of view was quantified from DAPI-stained images using a manually defined rectangular area (130µm×200µm for Achilles’ insertion site and 60µm×130µm for Supraspinatus) at the mid enthesis region (Figure 6). Selected regions were verified by brightfield images of the insertion sites. Images were converted to greyscale, thresholded, and pre-processed. Particle analysis was subsequently performed using ImageJ software.^46^

#### Immunohistochemistry

Expression of type X collagen (Col X) is an indicator of chondrocyte hypertrophy and marks the mineralization front of both the growth plate and enthesis.^3^ Immunohistochemistry was used to assess the distribution of Col X, marker of the mineralization front in the ECM. Sectioned slides from young samples were deparaffinized and rehydrated to 70% ethanol (n >= 4/genotype). Heat-mediated antigen retrieval was performed at 65°C (sodium citrate Buffer, pH 6.0). Slides were quenched and blocked at room temperature using 0.3% hydrogen peroxide (Santa Cruz, Dallas, TX) and 5% goat serum in PBS, respectively. Primary rabbit monoclonal anti-collagen X antibody (Abcam, ab260040; 1:100) with HRP/DAB system (Millipore Sigma) was used for detection of Col X. Slides were counterstained with Hematoxylin and cover slipped with acrylic mounting media (Acrymount, Statlab, McKinney, TX, USA). Presence of Col X in the ECM was quantified at the secondary ossification center by segmentation based on cellular and tissue morphology, and the region with Col X localization was measured using ImageJ software.^46^

### Mechanical testing

For biomechanics, frozen contralateral hindlimbs were thawed overnight and calcanei were dissected with minimal interruption of the Achilles attachment site. Bone-tendon complexes were equilibrated in PBS at room temperature prior to testing. Plantaris tendon and the gastrocnemius/soleus muscles were carefully removed without disruption of the Achilles tendon and enthesis. To avoid slip, samples were clamped in a custom-made fixture using sandpaper. Mechanical tests were performed using an electromechanical uniaxial tester (Instron 5943, Norwood, MA). All samples were tested in a saline bath at room temperature. For each sample, the major diameter area and gauge length were measured at 0.01N preload using a scaled image captured on video in frame of the preloaded sample. The cross-sectional area was then calculated with the assumption of ellipsoidal geometry with a diametric ratio of 7/5. The loading protocol consisted of a ramp to 0.02N, preconditioning with 10 cycles (0.02-0.04N), and displacement to failure at 0.03mm/s rate. Force-displacement data were collected and analyzed using a custom Matlab code to calculate stiffness, maximum load, work to maximum load, elastic modulus, ultimate stress, strain at ultimate stress, and area under the curve (i.e., toughness).

### Statistical analysis

Statistical analyses were performed using Prism (Graphpad, La Jolla, CA). Quantitative data are presented as dot plots with mean +/- 95% confidence interval unless otherwise indicated, and male and female mice are annotated as blue and pink dots, respectively. Results from bone morphometry (microCT), nuclear and Col X quantification, and uniaxial tensile testing were compared between WT and *Fgf9*^*ScxCre*^ samples using two-tailed unpaired t-tests (assuming Gaussian distribution). Comparisons between mouse ages for cellularity measurements of neonatal, young-adult, and adult entheses, were compared using a one-way ANOVA.

## Acknowledgments

Thanks to Dr. Gwen Talham and Frank Warren for assistance with animal care at the University of Delaware and to Zachary Tata for assistance with animal care at the University of Michigan. Funding for this work was provided by the National Institutes of Health (R01AR079367 and R03HD094594 to MLK, with support from K12HD073945, P30GM103333, and P30AR069620), (R01AR079246 and R01HD049808 to DMO); the National Science Foundation (1944448 to MLK); the University of Delaware Research Foundation (16A01396 to MLK); and the University of Delaware Doctoral Fellowship (EG). Schematics generated in Biorender.

